# Detecting pathogen exposure during the non-symptomatic incubation period using physiological data

**DOI:** 10.1101/218818

**Authors:** Lauren Milechin, Shakti Davis, Tejash Patel, Mark Hernandez, Greg Ciccarelli, Steven Schwartz, Siddharth Samsi, Lisa Hensley, Arthur Goff, John Trefry, Sara Johnston, Bret Purcell, Catherine Cabrera, Jack Fleischman, Albert Reuther, Franco Rossi, Anna Honko, William Pratt, Albert Swiston

**Affiliations:** Massachusetts Institute of Technology Lincoln Laboratory, Lexington MA, USA; US Army Medical Research Institute of Infectious Diseases, Ft. Detrick MD, USA

## Abstract

Early pathogen exposure detection allows better patient care and faster implementation of public health measures (patient isolation, contact tracing). Existing exposure detection most frequently relies on overt clinical symptoms, namely fever, during the infectious prodromal period. We have developed a robust machine learning based method to better detect asymptomatic states during the incubation period using subtle, sub-clinical physiological markers. Starting with high-resolution physiological waveform data from non-human primate studies of viral (Ebola, Marburg, Lassa, and Nipah viruses) and bacterial (*Y. pestis*) exposure, we processed the data to reduce short-term variability and normalize diurnal variations, then provided these to a supervised random forest classification algorithm and post-classifier declaration logic step to reduce false alarms. In most subjects detection is achieved well before the onset of fever; subject cross-validation across exposure studies (varying viruses, exposure routes, animal species, and target dose) lead to 51h mean early detection (at 0.93 area under the receiver-operating characteristic curve [AUCROC]). Evaluating the algorithm against entirely independent datasets for Lassa, Nipah, and *Y. pestis* exposures un-used in algorithm training and development yields a mean 51h early warning time (at AUCROC=0.95). We discuss which physiological indicators are most informative for early detection and options for extending this capability to limited datasets such as those available from wearable, non-invasive, ECG-based sensors.

## Introduction

We have developed a method for assessing pathogen exposure based solely on host physiological waveforms, in contrast to conventional diagnostics based on fever or biomolecules [1] of the pathogen itself or the host’s immune response. Early warning of pathogen exposure has many advantages: earlier patient care increases the probability of a positive prognosis [2-5] and faster public health measure deployment, such as patient isolation and contact tracing [6-8], which reduces transmission [9]. Following pathogen exposure, there exists an incubation phase where overt clinical symptoms are not yet present [10]. This incubation phase can vary from days to years depending on the virus [11, 12], and is reported to be 3-25 days for many hemorrhagic fevers [3, 4, 13, 14] and 2-4 days for *Y. pestis* [15]. Following this incubation phase, the prodromal period is marked by non-specific symptoms such as fever, rash, loss of appetite, and hypersomnia [10]. Fig 1 presents a conceptual model of the probability of infection detection *P_d_* during different post-exposure periods (incubation, prodrome, and virus-specific symptoms) for current specific and non-specific (i.e., symptoms-based) diagnostics. We also include what may be considered an “ideal” sensor system capable of detecting pathogen exposure even during the earliest moments of the incubation period. We hypothesized that quantifiable abnormalities (versus a diurnal baseline, for instance) in high-resolution physiological waveforms, such as those from electrocardiography, hemodynamics, and temperature, *before* overt clinical signs could be a basis for the ideal signal in Fig 1, thereby providing advanced notice (the early warning time, *Δt=t_fever_-t_ideal_*) of on-coming pathogen-induced illness.

**Fig 1:**
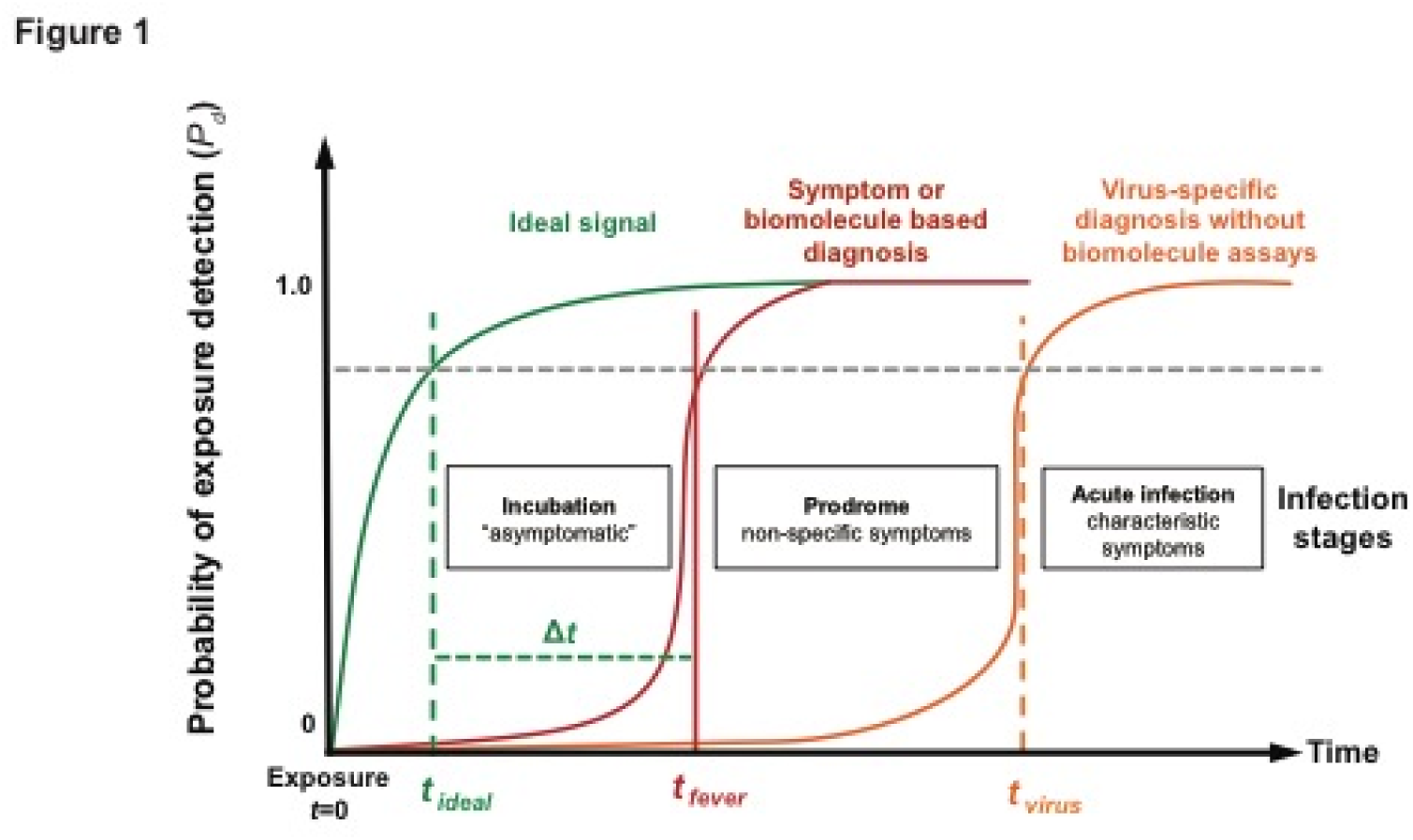
Phases following pathogen exposure. This notional schematic shows the probability of detection (*P_d_*) for current symptoms-based detection (red curve) and an ideal signal (green curve) versus time (viral exposure at *t*=0), overlaid with a typical evolution of symptoms. An ideal sensor and analysis system would be capable of detecting exposure for a given *P_d_* (and probability of false alarm, *P_fa_*) soon after exposure and during the incubation period (*t_ideal_*), well before the non-specific symptoms of the prodrome (*t_fever_*). We define the difference Δ*t* = *t_fever_ - t_ideal_* as the *early warning time*.

In addition to characteristic clinical presentations, most infectious disease diagnosis is based upon identification of pathogen-specific molecular signatures (via culture, PCR/RT-PCR or sequencing for DNA or RNA, or immunocapture assays for antigen or antibody) in a relevant biological fluid [10, 16-23]. Exciting new approaches allowed by high-throughput sequencing have shown the promise of pre-symptomatic detection using genomic [24, 25] or transcriptional [26-28] expression profiles in the host [29]. However, these approaches suffer from often prohibitively steep logistic burdens and associated costs (cold chain storage, equipment requirements, qualified operators, serial sampling): indeed, most infections presented clinically are never definitively determined etiologically, much less serially sampled. Furthermore, molecular diagnostics are rarely used until patient self-reporting and presentation of overt clinical symptoms, such as fever. Past physiological signal-based early infection detection work has been heavily focused on systemic bacterial infection [30-35], and largely centered upon higher sampling rates of body core temperature [35, 36], advanced analyses of strongly-confounded signals such as heart rate variability [31-33] or social dynamics [37], or sensor data fusion from already symptomatic (febrile) individuals [38]. While great progress has been made in developing techniques for signal-based early warning of bacterial infections and other critical illnesses in a hospital setting [39-42], we are aware of only one prior effort to extend these techniques to viral infections or other communicable pathogens in non-clinical contexts using wearable sensor systems [43].

Electronics miniaturization has led to a wave of wearable sensing technologies for health monitoring [44], and increasingly more processing power is available to consumers to make meaningful use of these collected data [45]. Inspired by these developments, we envision a low ergonomic profile, robust, wearable, personalized and multi-modal physiological monitoring system persistently measuring signals capable of sensitive pathogen exposure and infection detection; here we present a pilot investigation into building algorithms that enable this vision. Such a system could cue the use of highly specific (but expensive) diagnostic tests, prompt low-regret responses such as patient isolation and observation, or advise clinicians of fulminant complications in already compromised patients. In future, possibly etiologically-specific iterations of this approach, knowledge of causative pathogens could inform very early therapeutic intervention. Furthermore, using very feature-limited datasets, such as those that could be collected using wearable sensor platforms, would enable this technique to be implemented in non-ideal clinical, athletic, and military environments. Transitioning this technology to these contexts is the focus of ongoing work.

## Results

In this pilot study, we use high-resolution (both fast sampling rates and finely quantized amplitudes) physiological data from non-human primates (NHPs) exposed via intramuscular (IM), aerosol, or intratracheal routes to one of several viral hemorrhagic fevers (Ebola virus [EBOV], Marburg virus [MARV], Lassa virus [LASV]), Nipah virus (NiV), or one bacterial pathogen (*Y. pestis*) to build a high sensitivity, low etiological specificity (i.e., not informative of particular pathogens) processing and detection algorithm (see Fig 2a). Physiological data is standardized to remove diurnal rhythms, aggregated to reduce short-term fluctuations, and then provided to a supervised binary classification (exposed and unexposed classes) machine learning algorithm (Fig 2b). Supervised machine learning algorithms learn data characteristics that belong to pre-determined classes, then place new, unseen data into the appropriate class based on similar characteristics. Here, we define pre- and post-exposure as the two classes since “infection” itself is not a discrete event and all exposures in these studies lead to infection and illness. We tested and compared several classifiers; random forests had the best positive predictive value (discussed below) and were chosen for the rest of our analysis [46]. Random forests were also chosen for their high classification accuracy, robustness to many correlated features, and the ability to estimate the importance of variables in classification [46, 47]. We chose to grow (train) random forests at two post-exposure stages, thus allowing the algorithms to adapt to physiological changes between incubation and prodromal phases: one random forest is trained using post-exposure but pre-fever physiological data, and the other using post-exposure, post-fever data. Both random forest training sets include pre-exposure data to build the unexposed class. For algorithm evaluation, subject data is separated into various training and testing sets, and every testing subject’s data is provided to the random forest model for an exposure prediction every 30 min. After using binary integration and a constant false alarm thresholding approach to further reduce false alarms (Fig 2c), mean exposure declaration times are found to range from 32.6±40.5h (for LASV) to 74±37h (for NiV) before the onset of fever (defined as 1.5ºC above a diurnal baseline [48] sustained for two hours). We note that once the random forests have been trained, all physiological data is given to both pre- and post-fever models, without regard to exposure or fever status; in other words, our approach does not require information on exposure or fever times for successful classification and detection. This approach allows for both flexible, multi-modal input features (customizable to the available sensing hardware) and tunable false alarm rates, which offers a unique ability to adjust system performance per user needs. Additionally, our method leverages supervised classification to learn subtle physiological changes, and continuously monitors for signs of pathogen exposure rather than relying on a single time ‘snapshot’ of subject data.

**Fig 2:**
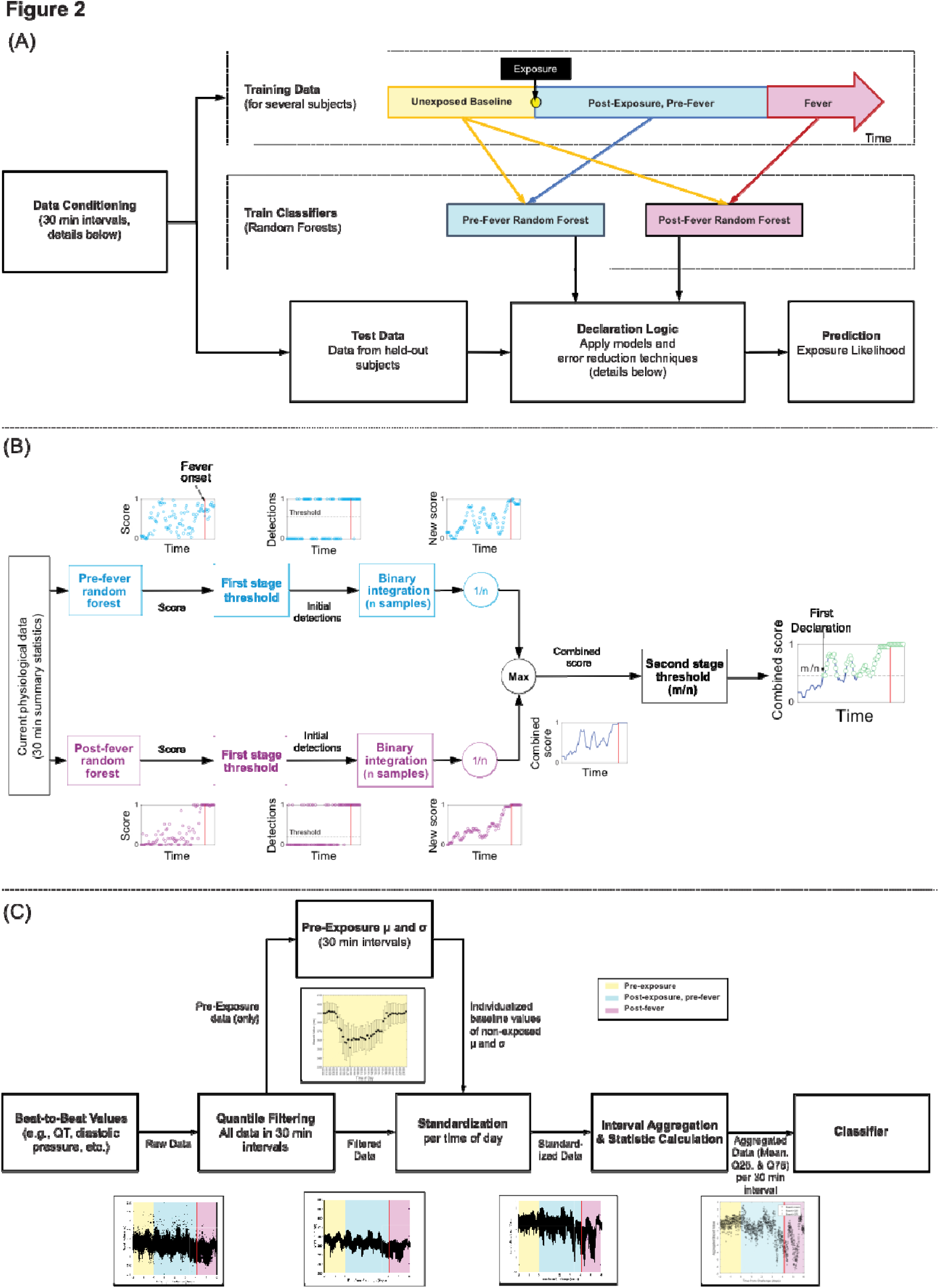
Overview workflows of our early warning algorithm. (a) Overview of our classification approach using random forests, including which data comprises the training sets for the two random forest classifiers. (b) Detail on how beat-by-beat data is conditioned to remove noise and diurnal cycles, as well as summary statistics computation which are the features provided to the two classifiers. (c) A block diagram of a two-stage detection algorithm to reduce false alarms. The detection scheme comprises two distinct stages: after the random forest model score output, an *a priori* determined threshold (based on a desired *P_fa_*) is applied to yield initial detections. These are then subjected to a binary integration step of the past *n* samples, and the maximum value of the pre- and post-fever models is taken to produce a single time series. A second stage *m* of *n* detection is applied, which finally produces a final ‘declaration’ of being exposed or not. All example data shown in (b) and (c) are from subjects in the MARV aerosol study. See Methods for detailed descriptions.

### Data Preprocessing and Detection Algorithm

Before classification, several data processing steps are required to remove time as an implicit feature in our physiological datasets (see Fig 2b). First, data is standardized and aggregated subject-by-subject to eliminate short-term fluctuations and daily diurnal rhythms. From these standardized datasets, mean and quantiles are calculated for each time window; these statistical measures are the features provided to the machine learning algorithm (see S2 Table for a list of features considered, and S10 Data for the complete dataset). Windows of length 30 minutes were chosen as a tradeoff between computational requirements and algorithm performance (as indicated by random forest out-of-bag errors; see S3 Fig for results using different length windows). For the rest of our analysis, data from 12h before and 24h after viral or bacterial challenge are excluded from performance metrics due to differences in animal handling and exposure sedation that resulted in significant physiological deviations from baseline data unrelated to pathogen infection (as seen in Figs 4 and 5 during the “Excluded” zone). Further details on data processing may be found in the Methods section.

After data is standardized and aggregated, these features are used to train a random forest classifier. This resultant ensemble is a collection of fifteen binary decision trees which then “vote” on whether given new data belongs in the exposed or unexposed class. In our work, more than fifteen trees in a random forest did not significantly decrease the out of bag error, which measures classification success (see S4 Fig for details). In our final model, two random forests are trained to detect the post-exposure class at distinct time epochs: one model is tuned to detect subtle markers during the incubation phase prior to fever, while the second model is tuned for the early prodromal phase (i.e., onset of overt febrile symptoms) where temperature-related features emerge as powerful discriminants. The training data for the pre-exposure class for both models is a subset of baseline data prior to challenge and the quantity of training data has been balanced for the negative (pre-exposure) and positive (post-exposure) classes to avoid biasing one class over the other. To select the ideal features to put in these final forests, we inspect the feature importance metrics given by random forests built consecutively on a reducing feature set. In this way, we pick the top ten features selected by results from a cross-validation set (see S4 Fig for details), and the final models are built with these features. The output of these random forest ensembles, however, is prone to false alarms, and we employ a two-stage detection logic process to reduce false positives to a pre-determined target level (we chose a target *P_fa_*=0.01). Final declarations of “exposed” or “unexposed” are the output of this two-stage process, and are reported below; further details of this detection logic can be found in the Methods section.

We experimented with several classification methods, including Naïve Bayes [49], k-Nearest Neighbors [50], and random forests, and compared each across sensitivity, false alarms, and early warning time metrics. All classifiers had positive predictive values (results in S1 Fig), yet we chose random forests for several reasons. Most importantly, random forests do not assume statistical independence of features, which is useful given highly correlated physiological feature sets [51]. They also provide a quantitative feature importance metric which facilitates post-hoc comparison to the known viral pathology sequence, thus providing mechanistic understanding of why these physiological anomalies are present, and which sensor types provide the most value. Furthermore, the most discriminating features can be selectively chosen to re-grow forests and allow for better algorithm performance with fewer feature inputs, helpful in addressing the dilemma of having many more features than samples or subjects producing them [52]. Finally, in empirical comparisons of many machine learning methods, random forests consistently rank among the best approaches [53], and we too found them to produce the best outputs among the classifiers tested. Though there are numerous other possible classifiers available, the purpose of this pilot study is to prove the concept of early detection rather than comparing possible learning methods; improved classification approaches are the subject of on-going work.

### Evaluation: Three-fold Cross-Validation

We developed and tested our machine learning approach with three initial exposure study datasets based on MARV IM, MARV aerosol, and EBOV aerosol exposures. Data from across all three studies are aggregated and used to train the random forest model, thus not requiring *a priori* individual baseline data to build the “not exposed” class. (An individual’s baseline data is still used to standardize features; see Figure 2b.) These models are then tested in a three-fold cross-validation scheme where each partition is composed of randomly-selected subjects from each of the three exposure studies (i.e., the group of subjects in a partition is *not* the same as a cohort in an exposure study; exact partition assignments can be found in S5 Table). In doing so, this explicitly varies 5 experimental variables (species and gender of animal, exposure route, pathogen, and target dose; see Table 1 below) across the three partitions, which reduces the likelihood of biasing the model for any particular condition. Algorithm performance for one representative subject (whose early warning time is closest to the studies’ mean) is shown in Fig 3a. The blue curve is the combined score output by the algorithm as a function of time, the red vertical line the fever time, and the green dashed vertical line denotes the first true positive declaration. Each green circle indicates a declaration, which if found before pathogen exposure, represents a false alarm. Therefore, the early warning time *Δt* is the time between green and red vertical lines. (However, while *Δt* is clinically very useful, we note that for our datasets the *mean* early warning time is an unstable performance metric since small changes in the number of subjects and detection logic thresholds can have large impacts on *Δt_mean_*.) In this cross-validation scenario, we find a system probability of detection *P_d_*=0.80±0.01 (i.e., correctly declaring a subject as being exposed after the pathogen challenge), a pre-fever *P_d_*=0.56±0.02, a system probability of false alarm *P_fa_* =0.013±0.003 (i.e., incorrectly declaring a subject as exposed before the pathogen exposure), and *Δt_mean_ =*51.0±11.9h based on 9931 decision points and *N*=20. While the value of the pre-fever *P_d_* may seem low, our definition considers all detection opportunities (i.e., every 30 min interval) in the pre-fever period, and does not reflect how detections evolve with time as the pathogen replicates and the host mounts an immune response. As such, we may expect few declarations of exposure immediately following the challenge, since the pathogen has not yet prompted a systemic response captured in the physiological data; as the host mounts an increasingly strong immune response, the likelihood of capturing this systemic response increases. (Fig 3c, described below, also reflects this time evolution of detection.) To add additional context to the *P_d_*, we present a measure of declaration confidence called “early warning purity,” which is a ratio of true positives to detection opportunities that occur between the first true positive declaration and before fever, for each subject in S5 Table. This measure captures the reliability of a declaration, where values closest to 1.0 indicate no false negatives after the first declaration.

**Fig 3:**
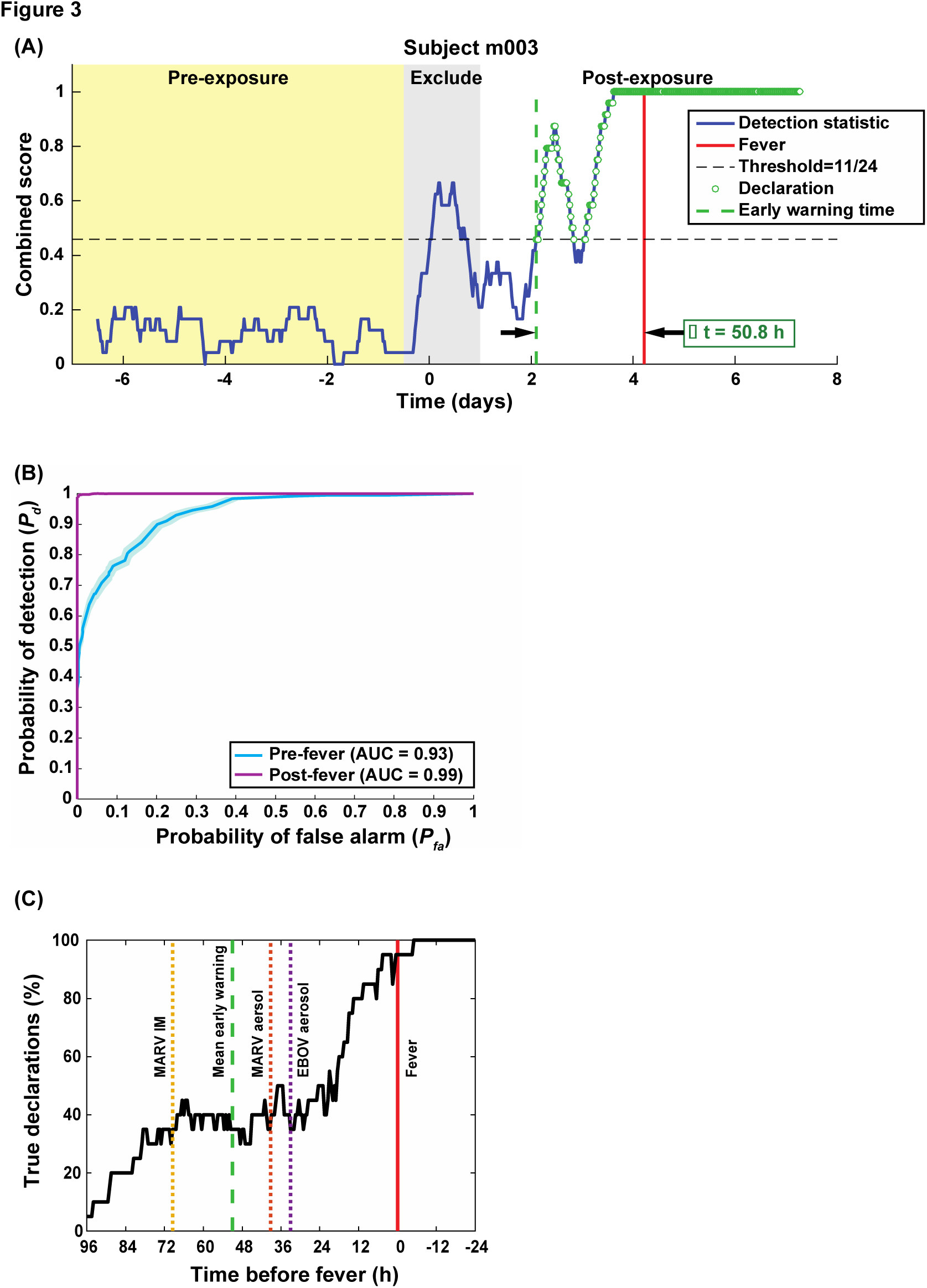
Algorithm output and performance measures from the three-fold cross-validation. (A) shows the combined score vs. time for a representative subject taken from the MARV aerosol exposure study, as well as declarations (green circle overlays) made by our detection threshold and binary integration. The combined score remains below the detection threshold (dashed horizontal line) before virus challenge, rises sharply around exposure (which is excluded) due to anesthesia, then rises again at ~2 days post-exposure when we make the first “exposed” declaration (dashed vertical green line). Combined score values below the detection threshold after exposure represent false negatives and the time between the first declaration and fever is this subject’s early warning time Δ*t*. The declaration purity in this early warning period is 91%. (B) presents the ROC curve, indicating nearly perfect performance after febrile symptoms, and strong positive predictive power (AUCROC=0.93) before fever. (C) shows the algorithm sensitivity (True Declarations) vs Time before fever for all 20 subjects, as well as the mean Δ*t* (vertical dashed lines) for each of the three constituent studies. (All other performance metrics may be found in S8 Table.) We find that half of the subjects are correctly identified as exposed 24-36h before fever, regardless of the particular pathogen, exposure route, or target dose.

**Table 1:** Summary of NHP studies used. The EBOV study compared two different physiological monitoring systems but data was combined and treated identically.

| Pathogen | Exposure | Subjects | Species | Monitoring | Target |
| --- | --- | --- | --- | --- | --- |

|  | method | (male/female) |  | system | dose |
| --- | --- | --- | --- | --- | --- |
| EBOV | Aerosol | 6 (3/3) | Cynomolgus | 3 subjects with ITS T37F<br>3 subjects with DSI L11 | 100 pfu |
| MARV [78] | Aerosol | 5 (3/2) | Rhesus | ITS T27F | 1000 pfu |
| MARV | IM | 9 (7/2) | Cynomolgus | ITS T27F | 1000 pfu |
| NiV [77] | IT | 5 (5/0) | African green monkey | ITS T27F | 20000 pfu |
| LASV [27] | Aerosol | 4 (4/0) | Cynomolgus | ITS T27F | 1000 pfu |
| *Y. pestis* | Aerosol | 4 (4/0) | African green monkey | ITS T27F | 100 LD<sub>50</sub> |

We further evaluated algorithm performance for all subjects by characterizing the system *P_d_* versus *P_fa_*, known as a receiver operating characteristic (ROC) curve (see Fig 3b)[54]. ROC curves describe the sensitivity (*P_d_*) and specificity (1-*P_fa_*, i.e., not informative of the causative agent) of a test and can be partially summarized by the area under the curve (AUC, where AUC=1.0 refers to a perfectly sensitive and specific detector, and AUC=0.5 indicates a test no better than a coin-flip). For this three-fold cross-validation, we find AUC = 0.93 for the pre-fever model, and AUC = 0.99 for the post-fever model, indicating strong positive predictive value during the “non-symptomatic” incubation period (where early warning is most meaningful) and nearly perfect performance during the symptomatic, febrile prodrome. This extremely high performance can be understood by considering that the post-fever classifier will dominate after febrile symptoms since elevated core body temperature is such a clear data feature with strong predictive power (and is how pathogen exposure assessments are most frequently performed). The final metric for algorithm performance we consider is shown in Fig 3c, which plots the percentage of subjects correctly declared as “exposed” (true positives) vs. early warning time, and is a measure of algorithm declaration sensitivity as a function of time given a target *P_fa_*=0.01. Each individual exposure cohort is shown as a dashed vertical line, which indicates individual differences between pathogens (and exposure study conditions). Within these three studies, we see earliest mean warning time for MARV IM exposure at *Δt_mean_*=69h, and the two aerosol exposures, EBOV and MARV, have similar mean values at *Δt_mean_*=33h and *Δt_mean_*=39h, respectively.

An additional output of the random forest models is an easily understood measure of relative feature importance; that is, which features provide the most accurate separation between exposed and non-exposed classes. The most discriminating features for the pre- and post-fever random forest models are identified from a set comprised of four feature types derived from temperature, ECG, blood pressure, and respiration measurements. (See S6 Table for a complete listing of most discriminating features in each model partition.) The random forest model reports features that follow clinical symptomology, namely that core temperature-based features (mean and quantiles of temperature) in the post-fever, prodrome model are the highest ranking in importance. Before fever, however, subtle ECG, blood pressure, and temperature derived features seem to be the highest ranking in feature importance, as has been reported at the earliest stages of sepsis [31-34] (see Discussion below). Among the hemodynamic features, quantiles of systolic and diastolic aortic pressure are among the most important. Among ECG-derived features, means and quantiles of QT intervals (corrected [55, 56] or not), RR intervals (inverse of instantaneous heart rate), and PR intervals are routinely selected as those with the greatest predictive capability. That both inter-and intra-cardiac cycle features are selected, and that the statistical distributions (rather than just the means) of ECG-based features emphasizes the value of high sampling rate waveform analysis, rather than single time point (such as Korotkoff sound based blood pressure) or averaged (heart rate based on observed beats per unit time) measures. Fortunately, ECG and temperature-based features are among the most consistent predictors throughout the six studies considered (since some studies used different monitoring hardware or software configurations), and allow us to apply these random forest models beyond the exposure studies used to train them.

### Evaluation: Testing on Independent Datasets

We further tested our algorithm’s extensibility for handling entirely independent data unavailable during model training and development. Whereas in the three-fold cross-validations above, models are tested on a held-out *subset* of data from within the same exposure studies, models can be also be trained on exposure study datasets and then be tested against *entirely independent datasets*. These new datasets were collected during studies using different pathogens, animal species, target doses, and exposure routes, just as above, and were collected in separate experimental protocols by different researchers at different times. To perform this type of validation, we train the random forest models using all subject data from the MARV IM, MARV aerosol, and EBOV aerosol studies from the previous section (including a mix of subject gender, see Table 1), then test the models against unseen data from LASV aerosol, NiV intratracheal, and *Y. pestis* aerosol exposures. Across all three pathogens, we find a system *P_d_*=0.90±0.007 and *P_fa_*=0.025±0.004, a pre-fever *P_d_* =0.55±0.03, and a *Δt_mean_*=51.0±13.9h. We show one representative subject for each pathogen in Fig 4. Even though the classifier was trained only on EBOV and MARV, we see significant pre-fever positive predictive value of the model, with an AUCROC=0.95 (Fig 5a). Fig 5a also shows the sensitivity vs. time curve for all subjects in the independent datasets, along with mean *Δt* for each pathogen exposure study. We find that NiV has the longest *Δt_mean_=*74h (though NiV subjects also have the longest incubation period, ~5days, and often these subjects have mediocre early warning purity values as seen in S5 Table), and that LASV and *Y. pestis* exposure studies have *Δt_mean_*=33h and *Δt_mean_*=41h, respectively (with a mean incubation period ~3.5 days).

**Fig 4:**
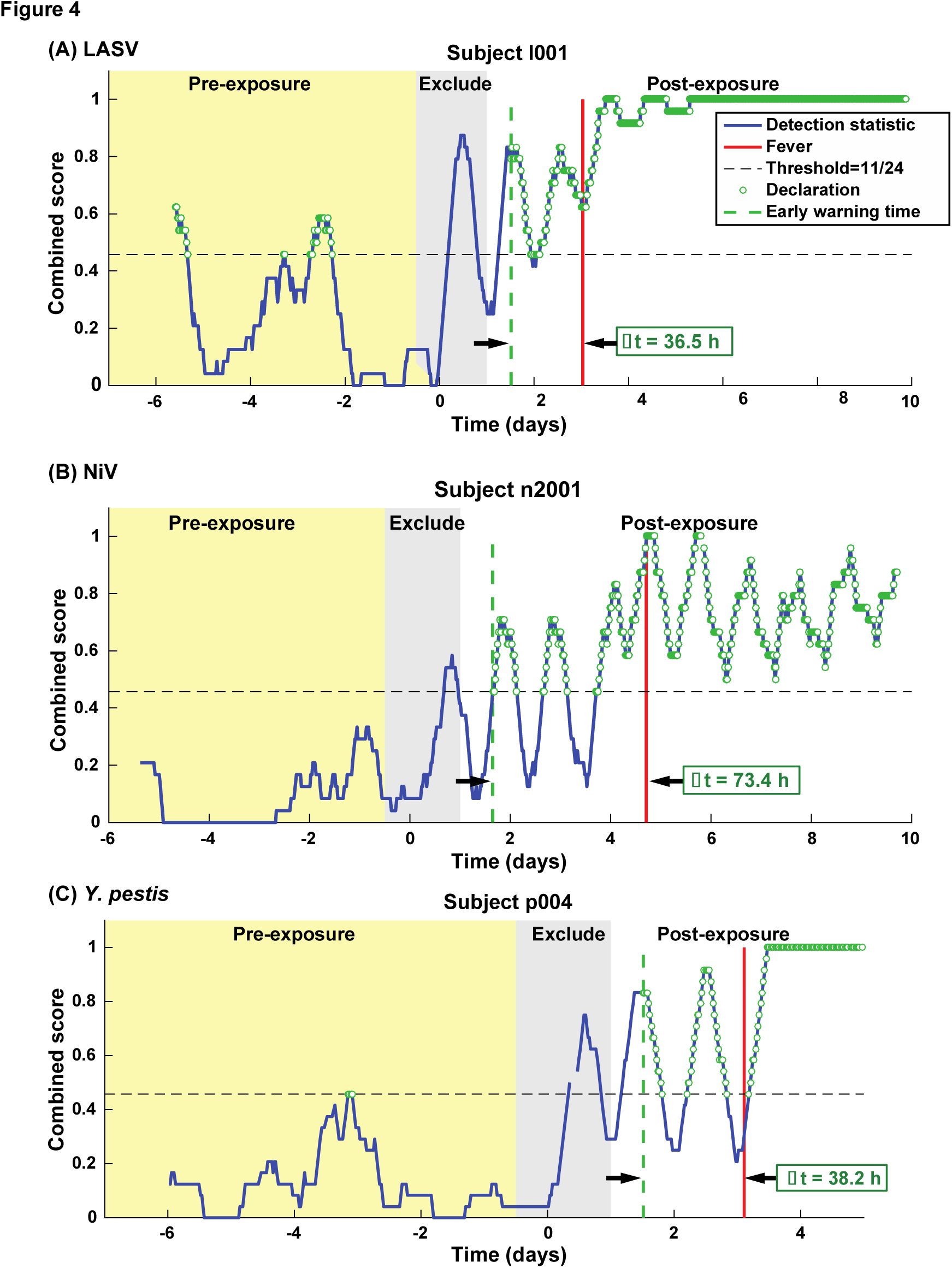
Representative single subject outputs from each of three independent datasets. Scores and declarations vs. time from the independent dataset validations for: (A) LASV, (B) NiV, and (C) *Y. pestis*. Declarations in the pre-exposure data represent false positives.

**Fig 5:**
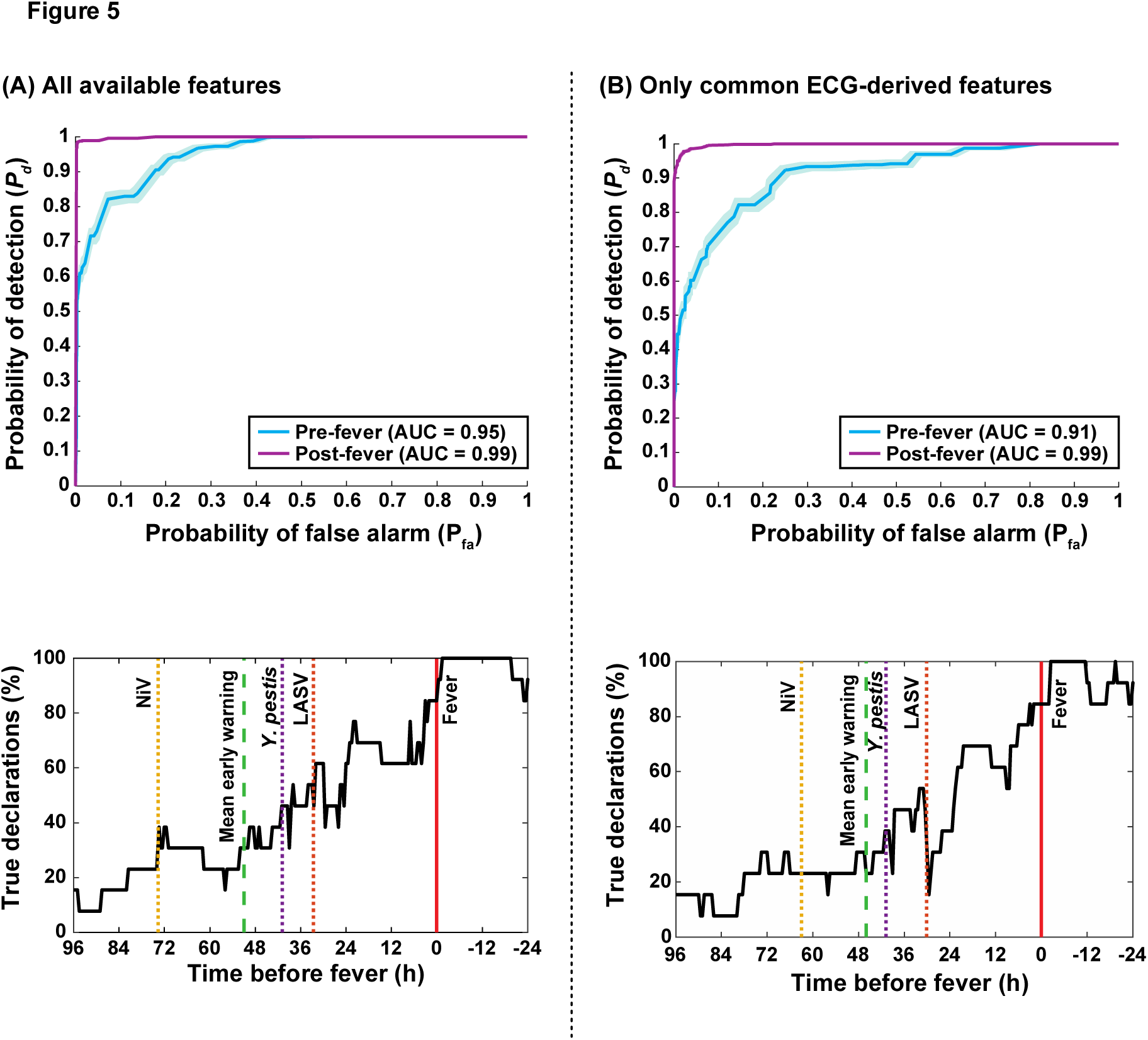
Algorithm output and performance measures from independent dataset validations. ROC and sensitivity vs. time before fever curves using (A) all available features from the implantable telemetry system, and (B) using only features derived from the ECG module that were common among all available studies (since some studies used different hardware or software configurations). Even when all temperature, hemodynamic, and pulmonary features are excluded, algorithm performance drops only slightly from Δ*t_mean_*=51h to 46h, and from pre-fever AUCROC=0.95 to 0.91. (All other performance parameters are available in S8 Table.) These results indicate that this type of early warning algorithm may possibly be embedded on an *ex vivo*, wearable ECG system such as a Holter monitor.

Unfortunately for the purpose of building early warning algorithm, the original animal exposure studies did not include “sham” animals to investigate the effects of exposure vehicle or animal handling. However, to build a synthetic “sham” dataset, we tested our algorithm with unexposed, pre-challenge subject data from the EBOV and NiV exposure studies that were otherwise excluded per the criteria given in the Methods section. These data include seven full days of measurements from each of nine animals prior to pathogen exposure: 7 subjects from the EBOV study (excluded due to therapeutic intervention following exposure) and 2 subjects from the NiV study (which developed fever earlier than our exclusion criteria). Detection results on these data from unexposed animals result in a consistently low false positive rate of *P_fa_*= 0.017±0.005.

Using these independent validation sets, we find that the random forest models trained on the original set of EBOV and MARV exposure studies continue to provide clinically useful early warning times with a manageable false alarm rate even against pathogens, exposure routes, or animal species that were unavailable during training. This successful extension of an early warning classifier trained on EBOV and MARV for a hemorrhagic fever virus (LASV), a henipavirus (NiV), and a gram-negative coccobacillus (*Y. pestis*) suggests algorithm insensitivity to particular pathogens, and possible generalization for novel or emerging agents for which data has not or can not be collected.

### Extending to non-invasive monitoring platforms

Physiological data features provided to our algorithm were collected using surgically implanted monitoring devices; such data could never be expected from military service members, health care workers responding to an outbreak, hospital patients, or the general public. As an *in silico* simulation for limiting our dataset to what may be collected using a wearable monitoring device, we reduced the considered feature set to include only ECG-derived features such as RR, QT, QRS, and PR intervals. Successful use of ECG data as a predictor of physiological compensatory potential during shock has been reported [31, 32], and ambulatory Holter monitor devices collect exactly this type of data [57], as do even less obtrusive devices for performance athletes. Fig 5 compares our algorithm’s performance using all available features (Fig 5a) and features derived only from the ECG waveform (Fig 5b). We see only modest performance decreases in *Δt_mean_* (46.0±14.1h), pre-fever *P_d_* (0.55±0.03), and system *P_d_* and *P_fa_* (0.89±0.008 and 0.026±0.004, respectively), even though core temperature, and hence onset of febrile symptoms, is no longer an available feature. These results may be expected given the highly correlated nature of physiological data, but positively suggests the implementation of this algorithm with non-invasive, ECG-based monitoring equipment.

## Discussion

Non-biochemical detection of pathogen incubation periods using only physiological data presents an enabling new tool in infectious disease care. Previous work has shown that reducing transmission during the viral incubation period is as or more effective an intervention as reducing the inherent transmissibility (*R_0_*) of the pathogen in controlling emerging outbreaks [9]. However, there is no existing method to detect this non-symptomatic incubation period that is possibly extensible to mobile settings or wearable sensor systems, such as high-resolution ECG. We present initial results towards building a multi-modal, supervised machine learning algorithm capable of determining this incubation period using only physiological waveforms, based on data collected in NHPs infected with several pathogens. Using the random forest method we avoid over-fitting the models, demonstrated by successful testing and training on both different subsets of data within the same exposure studies, as well as testing on entirely independent exposure datasets. These cross-validations show the promise of extending this approach beyond a given animal model, exposure method, or virus. While we chose a target system *P_fa_*~0.01 that was supported by the limited subject numbers in the studies available, this would not lead to an acceptable daily false alarm rate of about one declaration every 2 days (for 30min windows). We estimate *P_fa_* should be ~10^-3^ or less, which corresponds to one false alarm approximately every 3 weeks of continuous monitoring (again, for 30min windows). Reducing this critical system parameter to more clinically acceptable levels is the subject of on-going work, and will require larger sample sizes or more refined processing algorithms. Furthermore, the effect of physiological confounders, such as intense exercise, arrhythmias, lifestyle diseases, and autochthonous or annual infections has not been explored in this initial study; only with promising early warning times across a range of conditions would these more complex studies be justified.

We postulate that immuno-biological events of the innate immune system – particularly systemic release of pro-inflammatory chemokines and cytokines from infected phagocytes [28, 58-62], as well as afferent signaling to the central nervous system [63, 64] – are recapitulated in hemodynamic, thermoregulatory, or cardiac signals which may be more easily measured and assessed than biomolecule markers for viral infection (via sequencing [24, 25, 29] or immunocapture approaches [16, 17]). For instance, prostaglandins (PG) are up-regulated upon infection (including EBOV [65, 66]) and intricately involved in the non-specific “sickness syndrome” [67]; the PGs are also known to be potent vascular mediators [68] and endogenous pyrogens [69, 70]. Recent work has shown how phagocytic immune cells (macrophages, in particular) directly modulate electrical activity of the heart [71]. Past work has clarified how tightly integrated, complex, and oscillating biological systems can become uncoupled [72-74] during trauma [75] or critical illness [34, 76] which would be captured in the comprehensive, multi-modal physiological datasets used in our present work. Finding that our algorithm provides early warning times for both viral and (albeit limited) bacterial exposures suggests that the “exposure signal” found by the random forest models arises from the innate immune system, and is a generalized indication of immune activation rather than a specific signal for particular pathogens. Rigorously pursuing this hypothesis would require additional high temporal resolution pathogen exposure datasets, including biochemical, immunological, neurological, and cardiovascular information. Transitioning this capability into clinical use will also require the controlled exposure and monitoring of human subjects, such as during periodic influenza, tetanus, or zoster vaccinations.

Previous work on genomic [24, 25] profiles of peripheral blood cells following acute influenza infection indicate specific host responses at just ~45h following exposure, corresponding to ~35h of early warning time. Our combined results suggest that the classic understanding of a “non”-symptomatic incubation phase may be incomplete: during viral incubation, subtle sub-clinical cues (genomic, transcriptional, and physiological) can be detectable with sufficiently high-sensitivity sensor and analysis systems. Better understanding of how biomolecular changes are captured in systemic physiological signals during pathogen infection would open further opportunities for better therapeutic administration both before and during infection, quarantine or isolation, and vaccine development.

Detecting pathogen exposure before self-reporting or overt clinical symptoms affords great opportunities in clinical care and public health measures. However, given the consequences of using some of these interventions and the lack of etiological agent specificity in our algorithm, we envision this current approach (after appropriate human testing) to be a trigger for ‘low-regret’ actions rather than necessarily guiding medical care. For instance, using our high sensitivity approach as an alert for limited high specificity confirmatory diagnostics, such as sequencing or PCR-based, could lead to considerable cost savings (an “alert-confirm” system). Public health response following a bioterrorism incident could also benefit from triaging those exposed from the “worried well.” Ongoing work focuses on adding enough causative agent specificity to discern between bacterial and viral pathogens; even this binary classification would be of use for front-line therapeutic or mass casualty uses. Eventually, we envision a system that could give real-time prognostic information, even before obvious illness, guiding patients and clinicians in diagnostic or therapeutic use with better time resolution than ever before.

## Methods

### Viruses

The Marburg Angola isolate used was United States Army Medical Research Institute of Infectious Diseases (USAMRIID) challenge stock “R17214” (Marburg virus/H.sapiens-tc/ANG/2005/Angola-1379c); this was used for both aerosol (rhesus macaques) and IM (cynomolgus macaques) studies. Cynomolgus macaques were exposed to Ebola virus/H.sapiens-tc/COD/1995/Kikwit-9510621 at a target dose of 100 pfu (7U EBOV; USAMRIID challenge stock “R4415”; GenBank # KT762962). African green monkeys were exposed to the Malaysian Strain of Nipah virus (isolated from a patient from the 1998-1999 outbreak in Malaysia, provided to USAMRIID by the Centers for Disease Control and Prevention). Cynomolgus macaques were exposed to the Josiah strain of the Lassa virus challenge stock “AIMS 17294” (GenBank #s JN650517.1, JN650518.1).

### Description of Animal Studies

Dr. William Pratt provided physiological data in NSS format (Notocord Systems, Croissy-sur-Seine, France) from adult (non-juvenile) non-human primate natural history studies previously conducted at the USAMRIID, summarized in Table 1. Research was conducted under an IACUC approved protocol in compliance with the Animal Welfare Act, PHS Policy, and other Federal statutes and regulations relating to animals and experiments involving animals. The facility where this research was conducted is accredited by the Association for Assessment and Accreditation of Laboratory Animal Care, International and adheres to principles stated in the Guide for the Care and Use of Laboratory Animals, National Research Council, 2011. Minimum number of subjects in MARV and EBOV studies was chosen using a Fisher exact test, with 100% lethality as the pre-specified effect, and thus all animals became infected following exposure. The study statistician randomized subjects for inclusion and pathogen exposure order by age, weight, and gender. No sham control subjects were included in the study design, and pre-exposure data was used to build the “un-exposed” class. In each study, remote telemetry devices (Konigsberg Instruments, Inc., T27F or T37F, or Data Sciences International Inc. L11: see details in Table 1) were implanted 3 to 5 months before exposure, and, if used, a central venous catheter was implanted 2 to 4 weeks before. NHPs were transferred into BSL4 containment 5 to 7 days before viral exposure, and baseline pre-exposed data collected for 4 to 6 days before. Subjects were exposed under sedation via either aerosol, intramuscular injection, or intratracheal exposure depending on the study, detailed in Table 1. The exposure time (*t*=0) used in our model is based upon the time of intramuscular injection or intratracheal exposure, or when a subject was returned to the cage following aerosol exposure (~20 min). All subjects were monitored until death or the completion of the study. Since these natural history studies involved no diagnostic tests or therapeutic interventions, and all subjects were administered infectious doses, there is no need for investigator blinding during the data collection phase. Since these natural history studies were not designed specifically for the purpose of algorithm development, no sham control animals were available, and all baseline unexposed and uninfected data were obtained from the pre-exposure period. Algorithm investigators were blinded to the study design until after animal data collection. The telemetry devices measure several raw physiological signals, which were translated to blood pressure (sampling frequency *f_s_* = 250Hz), ECG (*f_s_* = 500Hz), temperature (*f_s_* = 50Hz), and pulmonary (*f_s_* = 50Hz) features using Notocord software. We analyzed six separate exposure studies and used all subjects’ post-exposure data that had sufficient fidelity (i.e., no data loss from equipment failure), which developed fever two days or less before the studies’ mean (i.e., no possible co-morbid infections or complications), and did not receive a post-exposure therapeutic: these criteria led to 13 excluded animals, 2 from each the NiV and MARV IM studies, and 9 from the EBOV study (including 7 which received therapy). Some of the excluded EBOV and NiV subject’s pre-challenge data were used in the independent dataset validations to estimate thresholds (see below) and reduce the false alarm rate. Additional exposure study details for LASV, NiV, and MARV may be found elsewhere [27, 77, 78].

### Physiological Data Processing

All data processing and modeling was performed in Matlab (MathWorks, Natick MA). Physiological data is time dependent (that is, sequential time-series data) and is subject to short-term fluctuations and diurnal or circadian rhythms. Random forest classifiers, however, assume that the statistics of the data are independent of time and subject. To reduce diurnal and subject-to-subject dependencies from the data, each subject’s data is conditioned individually. The first conditioning step is to remove artifacts from motion, poor sensor placement or intermittent transmission drop outs by dividing the data into a series of *k*-minute intervals and omitting the top and bottom 2% quantiles for each interval. Next, we estimate baseline diurnal statistics for the *i*^th^ time-of-day interval during the pre-exposure period (*i.e*., data from several pre-exposure days, all corresponding to the same time of day, such as the thirty minute interval from 12:00PM to 12:30PM) by computing mean, *μ_i_*, and standard deviation, *σ_i_*. The data for the *i*^th^ time-of-day interval is standardized by subtracting the mean and dividing by the standard deviation from each data sample x_*i*_(*j*) in the corresponding *i^th^* interval, (*x_i_*(*j*) − *μ_i_*)/*σ_i_*. For a sufficiently short time interval of *k*-minutes, the data statistics are assumed to be approximately constant, therefore standardization mitigates diurnal time dependence from the signals. Then, three summary statistics are calculated for an *l*-minute block: mean and 25% and 75% quantiles. These time-independent summary statistics are the features for the random forest algorithm. We investigated the influence of values for *k* and *l* on successful classification, and while *k* and *l* do not need to be identical, chose *k*=*l*=30 min as a trade off between computational requirements and low random forest out-of-bag-errors (see S4 Fig). For example, *k*=*l*=30 min for two days of 4 raw physiological signals yields 96 time points with 12 data features. Data samples that correspond to measurements before pathogen challenge are labeled “0” to denote the pre-exposed class and those after challenge are labeled “1” to denote the post-exposure class.

### Random Forest Ensemble

Our model is composed of two random forests built using the TreeBagger class in the MATLAB Statistics and Machine Learning Toolbox. One random forest is grown using post-exposure training data prior to fever onset (labeled class “1”) and an equal number of randomly chosen negative data samples from the pre-exposure period (class “0”). The second random forest is trained similarly, but class “1” data corresponds to post-exposure training data after fever onset. Test data is always held out until the final evaluation step. Each random forest contains 15 classification decision trees grown on random subsets of data and features; 15 trees were chosen as a trade off between computational resources and successful classification, as indicated by the plateau in random forest out-of-bag-errors vs number of trees (see S4 Fig). The trees cast their “votes” for class “0” or “1,” and the forest returns a score equal to the proportion of trees that voted for the exposure (“1”) class. Random forests also provide feature importance metrics, and we use these metrics to find the most predictive features for difficult-to-classify pre-fever days. Initially all features are considered for training the random forest models, but once a subset of most predictive features is determined within a cross-validation training set, the random forest’s are regrown (on the original training dataset) using only the top 10 features to produce the final models upon which the corresponding testing set performance results are based. A rank order list of top 10 features from each study is provided in S6-S7 Tables.

### Detection Logic

We make declarations of exposure using a two-stage detection process (see Fig 2c). In stage one of the detection process, random forest model prediction scores (between 0 and 1 for every *l*=30 minute interval) are thresholded (i.e., a value of 1 is returned if the random forest model score is greater than or equal to a false alarm rate determined threshold, discussed below) to form a series of initial detections for the model every *l*=30 minutes. Threshold levels for both pre- and post-fever random forests are estimated by analyzing false alarm rates (Type I errors) of the initial detections versus threshold levels (swept from 0 to 1). The probability of false alarm (or *P_fa_*) is defined as

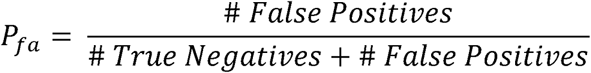

To enforce a desired significance level (we choose *P_fa_* = 0.01), we estimate the threshold needed to achieve a target *P_fa_* using a 3-fold approach similar to that used in random forest model training. For the case of validating performance on an independent test set (NiV, LASV, and *Y. pestis*), we randomly assign the test set subjects into 3 partitions for the purposes of threshold estimation. This approach maintains separation between the partition-under-test and the remaining two partitions used for threshold estimation, while providing a sufficient number of samples to estimate low rates of false alarms. Detections from the unexposed class of all but the partition-under-test are used to select the smallest first-stage thresholds (for pre- and post-fever as seen in Fig 2c) that support the desired *P_fa_*. This approach is repeated for each partition, resulting in independent estimates of the threshold pair (pre- and post-fever) for each partition. While a significance level of *P_fa_* = 0.01 is targeted, the overall system *P_fa_* may be higher or lower due to strict separation between the subjects-under-test and the subjects used to estimate the threshold.

These initial detections from each random forest model are subjected to a second-stage detection test to further reduce the false alarm rate. During the second stage, binary integration is performed over a sliding window of the past *n* initial detections. The accumulated detections are divided by *n*, giving a mean score for the pre- and post-fever random forest models. Next, scores are combined by taking the maximum of the pre- or post-fever values to create a single time series. At each 30 minute time interval, this combined score is compared to a final declaration threshold of *m*/*n*, where *m* ≤ *n* (we selected *n*=24 for a system latency of no more than 12 hours and selected *m*=11 which approximates the optimum binary integration threshold for a steady signal in noise [79]; performance is relatively insensitive to small deviations in *m* or *n*). The algorithm makes a ‘declaration’ that the subject is in the exposed class when the combined score is greater than or equal to *m*/*n*; if the threshold is not met, the algorithm assigns the subject to the ‘not exposed’ class for that time epoch. Note that *n* samples are required before a declaration can be made, so following the start of data collection or the end of an exclusion period (the 24h period following the challenge), no declarations are reported in the first *k*n* minutes (for *n*=24 and *k*=30min, this accumulation period effectively extends the exclusion period to 36 hours post-exposure).

### Model Performance Evaluation: Three-fold Cross-Validation and Independent Dataset Testing

Model performance may be evaluated by strictly separating subjects into testing and training sets. To characterize the performance, we conduct two modes of evaluation: *1)* an three-fold cross-validation, where a collection of exposure studies is used to develop and test the algorithms (data includes EBOV aerosol, MARV aerosol, MARV IM, and thus can vary in subject species, virus, and exposure route conditions), and *2)* an independent validation where models trained on the initial set of exposure studies (used in *(1)* above) are applied to an entirely new dataset with pathogens and experimental conditions not seen in the models’ training or tuning.

In the three-fold cross-validation mode of evaluation, subjects from the aggregated collection were randomly assigned into three partitions (each partition included animals from each of the 3 constituent exposure studies), which has been shown to perform better [80] than leave-one-out validations for smaller datasets. In turn, subjects from one partition were used to train the random forest models, the second partition was used as an independent cross-validation set to evaluate effects of tuning the model and algorithm parameters, and the third partition was used to evaluate final model performance. Model building and performance evaluation is repeated three times such that each partition is evaluated in each role. For mode *(2)* above, independent dataset testing is performed by treating all subjects from the three studies used in the initial set (EBOV, MARV IM and MARV aerosol) as a single training set to build and the random forest models and select the most important features. The resulting random forest models are then applied to previously unseen subjects from the LASV, NiV, and *Y. pestis* studies for the final performance analysis.

To evaluate system-level performance, we define probability of correct declaration *P_d_* as:

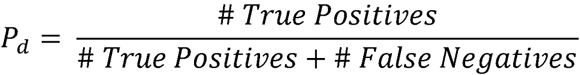

and *P_fa_* as above, where the True Positives, False Positives, True Negatives and False Negatives are evaluated on the final declaration outputs of Fig 2c. This definition includes all possible detection opportunities (every 30 minute interval), and does not retain any temporal information. To provide some temporal context, *P_d_* can also calculated for pre- and post-fever time epochs. In this case, the True Positives count is restricted to detection opportunities before and after fever, respectively. When reporting *P_d_* and *P_fa_* for a study and exposure condition, we include the 95% confidence interval based on normal distributions since the number of trials per study is large (>500 declaration points per class). Although some correlation is likely within a binary integration window of *k*n* minutes, we assume independence for trials separated by at least *k*n* minutes. We generate receiver operating characteristic (ROC) curves to measure system performance by calculating *P_d_* vs *P_fa_* at a series of threshold values (sweeping the first-stage detection threshold but holding the second-stage *m*/*n* threshold constant) and quantify the system performance with the ROC area under the curve (AUC), where an AUC=1.0 indicates perfect performance and AUC=0.5 indicates that the model is no better than a coin toss. Sensitivity (*P_d_*) is expected to be highest after febrile symptoms are apparent. To distinguish the sensitivity of the system during the pre- and post-fever epochs, *P_d_* is calculated independently before and after the onset of fever. The result is two ROC curves and corresponding AUCs: one evaluated on positive data restricted to pre-fever time samples and the other restricted to post-fever time samples. The negative data and two-stage detection process are identical for both ROC curves.

In a clinical early warning system, it may be desirable to calculate *P_d_* and *P_fa_* on a per-device, per-subject, or per-day basis. However, for this proof-of-concept study, the limited pool of subjects available (*N*=33 total) necessitates calculating *P_d_* and *P_fa_* across all 30-minute test points that are not in the exclusion window (12h before and 24h after exposure). This approach includes false negatives that may occur after an initial early warning declaration is made, and thus provides a conservative estimate of the device sensitivity which we predict will further increase with larger sample sizes and more refined processing algorithms.

Another important measure of system performance is the mean early warning time. The early warning time for an individual subject is defined as the time of the first true declaration (excluding data from the 24 h interval immediately following the challenge) minus the time of fever onset (defined as 1.5ºC above a diurnal baseline [48] sustained for two hours). Early warning times vary across subjects in a study, so the mean value is calculated across all subjects to characterize the early warning time afforded by the system. Since the number of trials (equal to the number of subjects) for this performance metric is relatively small, we bound the mean early warning time with a 95% confidence interval based on a *t*-distribution, and note that mean *Δt* is an unstable performance metric when evaluating small subsets of the data, such as on a per-pathogen level.

Model tuning, including feature selection and other classifier and detection parameters, may also be performed using an independent cross-validation testing set. A sweep over detection parameters *m* and *n* is provided in S9 Fig and illustrates some of the algorithm design trade-offs in selecting a short enough evaluation interval to allow for early warning while enforcing a long enough interval to maintain low false positives and high detection sensitivity prior to fever.

## Acknowledgements

We thank Jason Williams for his excellent graphics support, Dr. Brian Telfer for his thoughtful manuscript comments, and Amanda Casale for her expert statistics guidance. **Funding**. This material is based upon work supported by the Department of the Army and Defense Threat Reduction Agency under Air Force Contract #FA8721-05-C-0002 and/or FA8702-15-D-0001. Any opinions, findings, conclusions or recommendations expressed in this material are those of the author(s) and are not necessarily reflect the views of the United States Government or reflect the views or policies of the US Department of Health and Human Services. **Competing Financial Interests.** The authors declare competing financial interests: provisional U.S patent applications 62/193,961 and 62/337,964 were filed July 2015 and May 2016, respectively. **Data availability**. Data tables of standardized and aggregated physiology data, along with metadata, are available in the Supplementary Materials.

## Supplementary Materials Legends

S1 Fig: **ROC curves showing performance comparison of three different classifiers.** Random Forests, Naïve Bayes, and k-Nearest Neighbors methods were compared using the three-fold cross validation dataset. Shading around the ROC curves indicates 95% confidence intervals.

S2 Table: **Feature titles (prefixes and suffixes) and their descriptions.**

S3 Fig: **Determining optimal values for data processing parameters**. In the aggregated 3-fold cross-validation, sweeping values for the standardization (*k*) and aggregation (*l*) length versus RF out-of-bag error show a balance between computation intensity (higher for shorter windows) and RF classification errors. We chose *k*=*l*=30min (red box) as a compromise between these two system requirements. The total pre-classifier data processing time per subject per day on a Dell desktop computer (with two Intel Xeon E5607 processors and 12GB RAM) is significantly less than the individual standardization windows length itself.

S4 Fig: **Justification for choice of number of features (10) and trees (15) in our RF models**. The *P_fa_* (A&B) and *P_d_* (C&D) plateaus around 10 features for both pre- and post-fever models. Choosing the fewest features allows us to reduce processing time. (E) shows the RF out-of-bag-error plateaus around 15 trees used in the forest models. Choosing the fewest number of trees is one method to avoid over-fitting the model to the data.

S5 Table: **Declaration performance for each subject in all validation experiments**. Here, False Declarations refers to the number of false positives, Data Samples (unexposed class) is the sum of true negatives and false positives, True Declarations is the number of true positives, Data Samples (exposed class) is the sum of true positives and false negatives, and EW Purity is the ratio of True Positives to Data Samples between the first declaration and fever. The final rows detail model performance using only ECG-derived features in the independent dataset validations.

S6 Table: **Ranked feature importance for each partition in the aggregated three-fold validations before and after fever.**

S7 Table: **Ranked feature importance for the independent dataset validations before and after fever.**

S8 Table: **System performance metrics for all validations.** The aggregated three-fold cross-validation includes data from each of the three exposure studies in its training set. This same classifier is used to test independent LASV, NiV, and *Y. pestis* exposure study datasets including pre-exposure data from excluded subjects (see exclusion criteria under Description of Animal Studies subsection). The detection parameters for each study are *m*=11, *n*=24 and thresholds are estimated a priori for system *P_fa_* = 0.01. The broad distribution in *Δt* values both within and across pathogens can be understood both from the limited number of subjects for each pathogen (*N_LASV_* = *N_y.pestis_* =4 and *N_NiV_*=5) and different lengths of each pathogens incubation and onset of prodromal periods.

S9 Fig: **Performance evaluation across different detection logic parameters *m* and *n* for a target system *P_fa_* = 0.01.** The theoretical optimal value (see Ref 77 in manuscript) of *m* for a given *n* and *P_fa_* is indicated by the dashed line, and our operating point is indicated by the asterisk. (A) shows that small values of *n* promote earlier warning times (Δ*t*) by limiting the evaluation interval for a declaration. (b) The theoretical optimal value of *m* for a given *n* and *P_fa_* (dashed line) aligns with a relatively flat region of high *P_d_*. (c) shows that the actual system *P_fa_* is a few percent higher than the target system *P_fa_* of 0.01, but is relatively insensitive to the choice of *m* and *n* (except for very small ratios *m /n*). (d) Overall detection performance as measured by ROCAUC improves with larger values of *n*.

